## Supplementary Figures for "The evolution and convergence of mutation spectra across mammals"

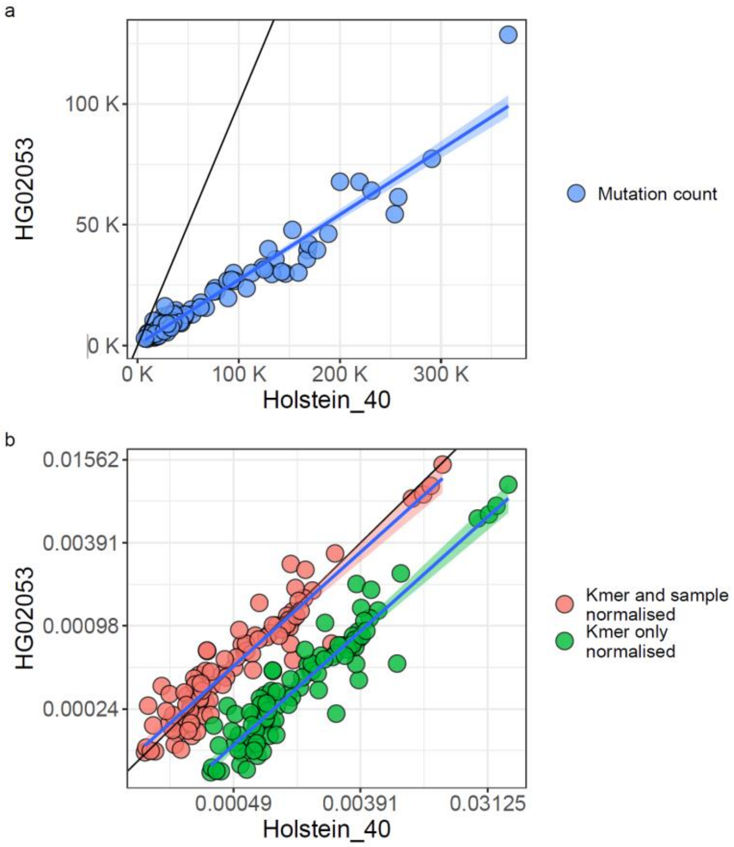


Supplementary Figure 1: Example impact of implemented normalisation approach. A) The raw mutation numbers in a randomly selected human (y axis) and cow (x axis) genome. Each dot represents the frequency of a given 3mer change. The parity line is shown in black. The cow genome carries substantially more mutations than the human genome. B) The impact of normalising by each genome's ancestral 3mer frequency (green) and the impact of further applying the median of ratios approach to each sample (red). Again the parity line is shown in black. Applying both corrections puts the mutation spectra of the two genomes on corresponding scales so that they can be more readily compared.

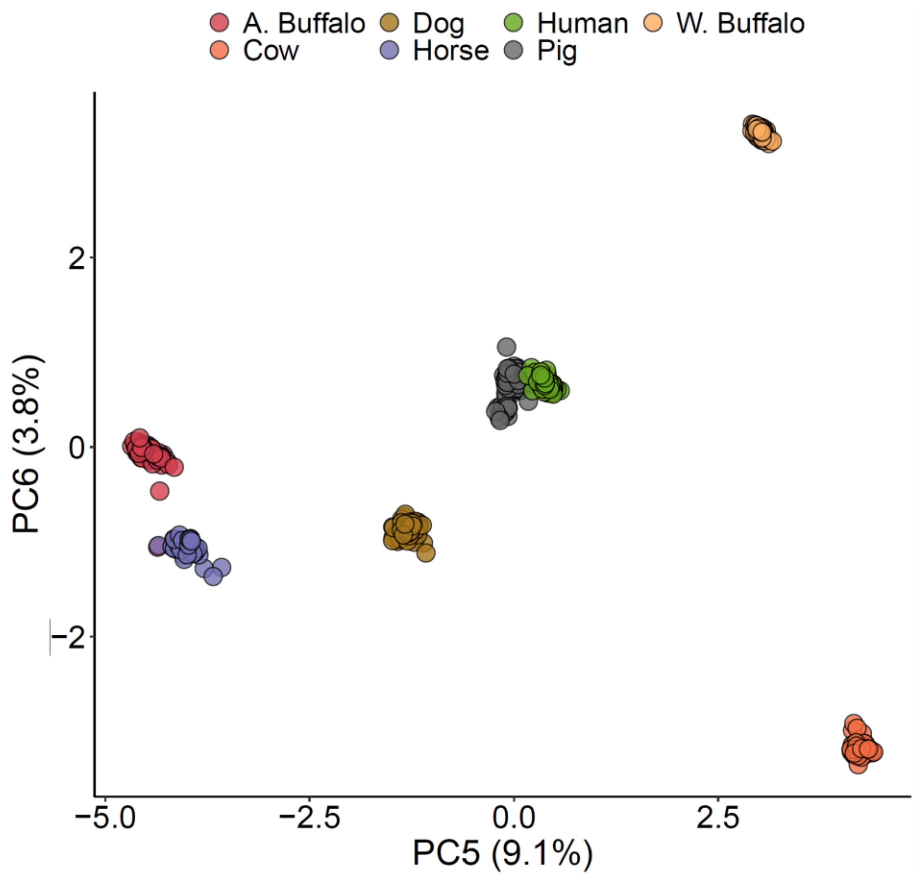


*Supplementary Figure 2. Principal component analysis of the relationship between different species based on the rate of SNV mutations of different ancestral 3-mers. PC5 vs PC6 is shown, illustrating how, despite their relatively close evolutionary distance, the Bovidae can be clearly separated by their underlying mutational spectra on these principal compenents.*

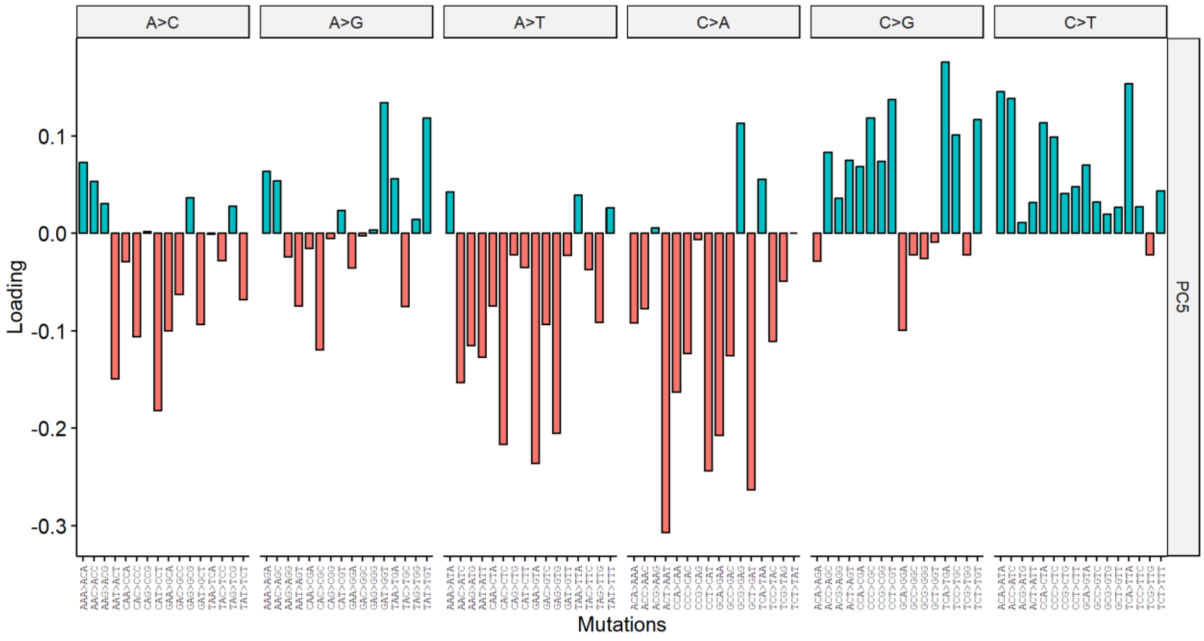


*Supplementary Figure 3. Loadings associated with PC5 in the between species SNV mutation profile comparison.*

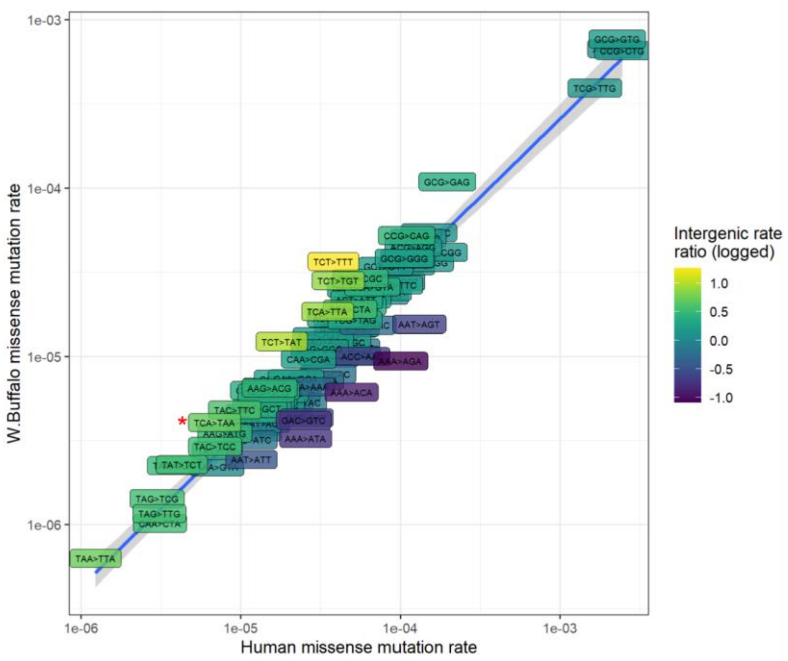


*Supplementary Figure 4. The rate of different changes that lead to amino acid changes in humans versus water buffalo. The colour of each point corresponds to the ratio of the rate of the same change between the same species, but in intergenic regions. The T[C>A]A change showing elevated rates in water buffalo is indicated.*

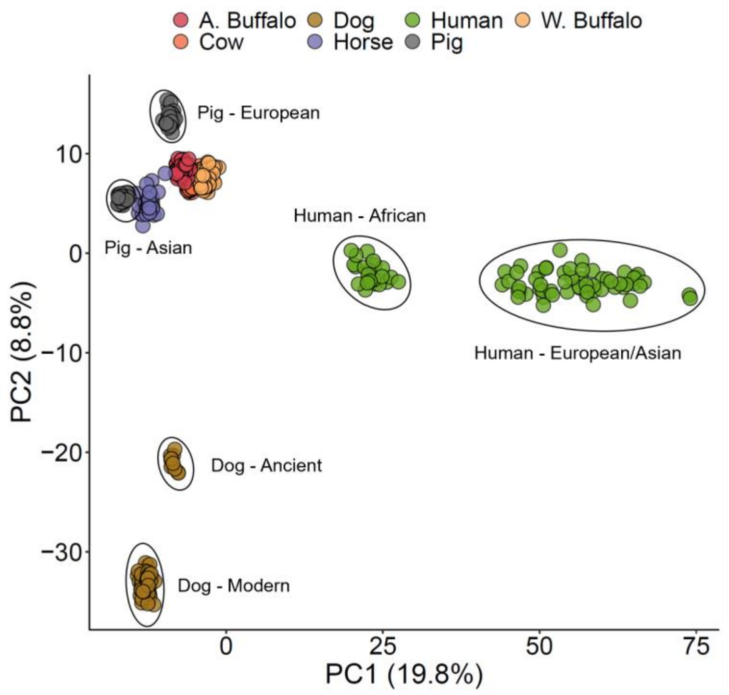


*Supplementary Figure 5. Principal component analysis of the relationship between different species based on the rate of SDM mutations of different ancestral 3-mers. The species with larger numbers were randomly downsampled to a maximum of 80 individuals. Further sub-division of the dog, pig and human populations is observed relative to the corresponding PCA for SNVs shown in Figure 2A.*

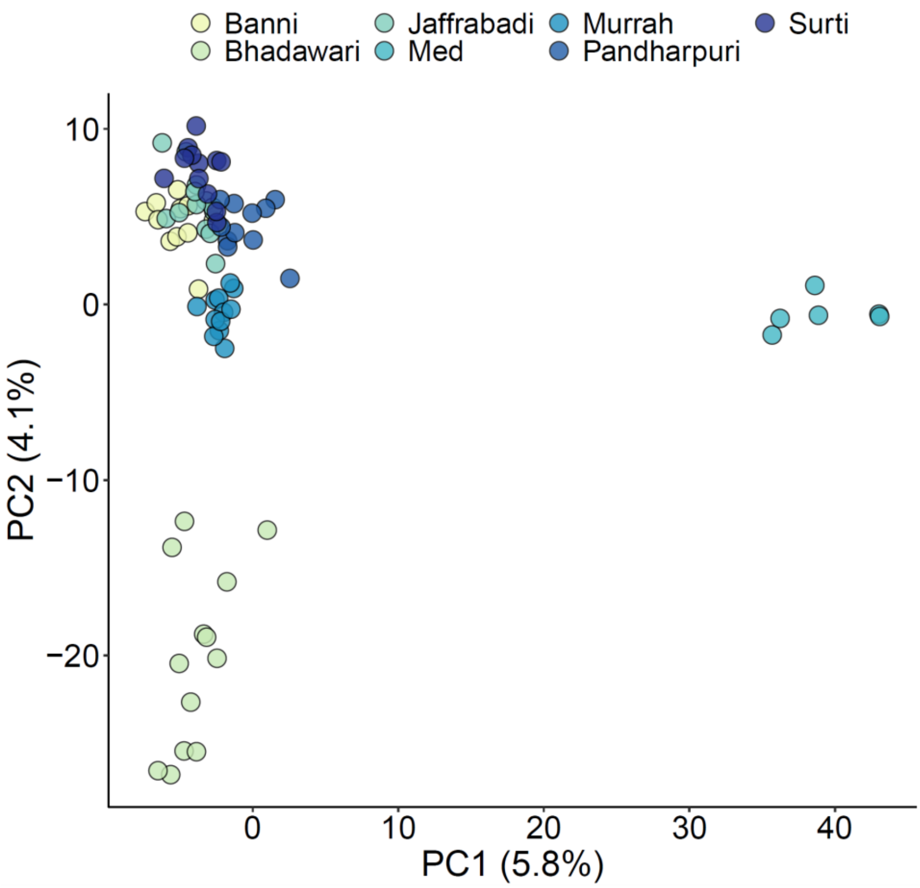


*Supplementary Figure 6. Principal component analysis of the relationship between different water buffalo breeds based on the rate of SDM mutations of different ancestral 3-mers.*

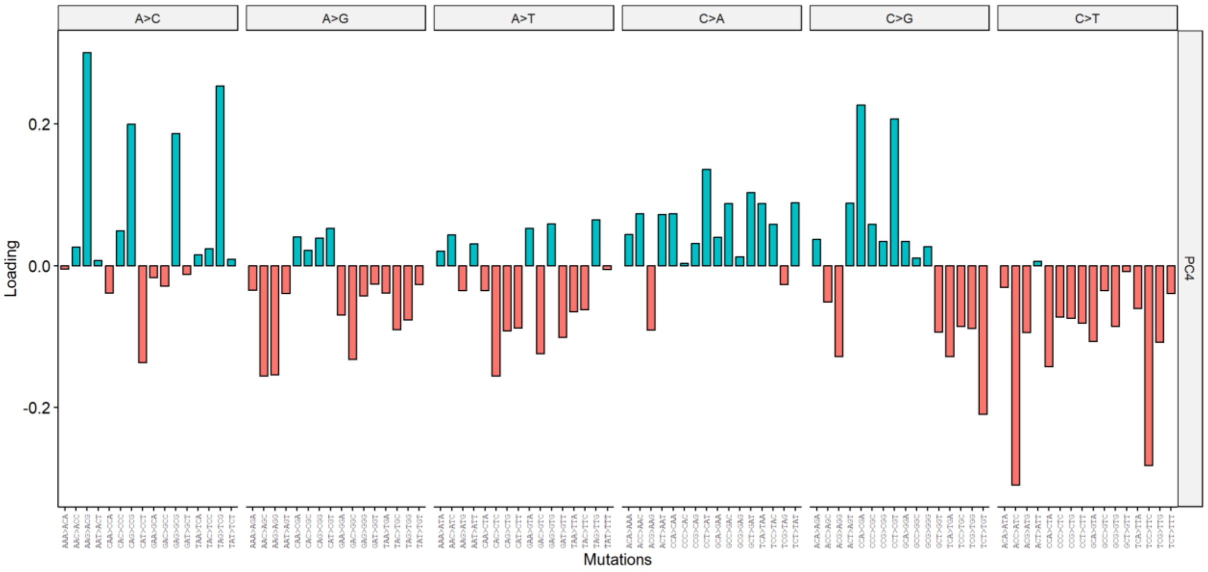


*Supplementary Figure 7. Loadings associated with PC4 in the between cattle population SNV mutation profile comparison. Positive loadings indicate a relative enrichment in African indicine populations. Negative loadings a relative enrichment in East Asian indicine.*

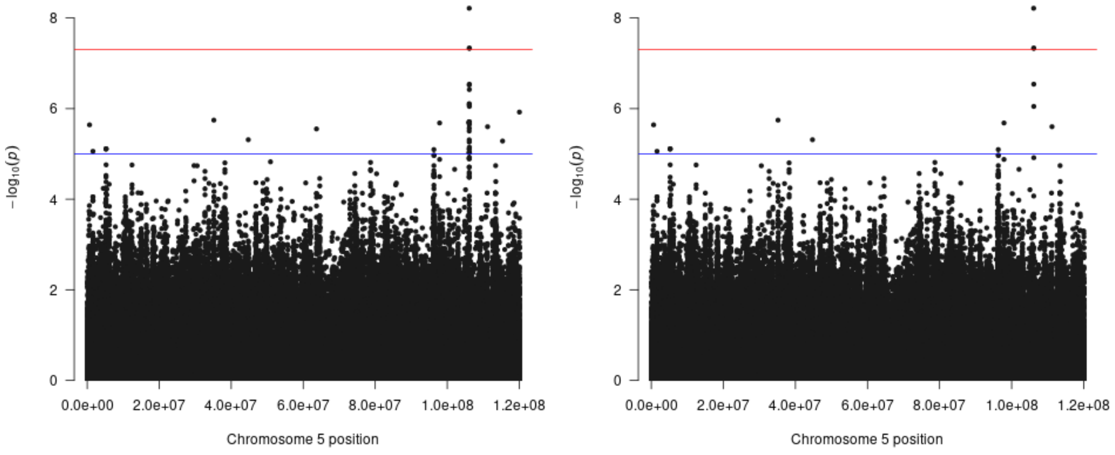


Supplementary Figure 8. PARP11 locus on chr5 with evidence of an association with the cattle indicine mutation profile captured by SNV PC4. Left figure is with a minor allele count cutoff of 20 (minor allele frequency of 0.067). Right figure is with a minor allele frequency cutoff of 0.1.

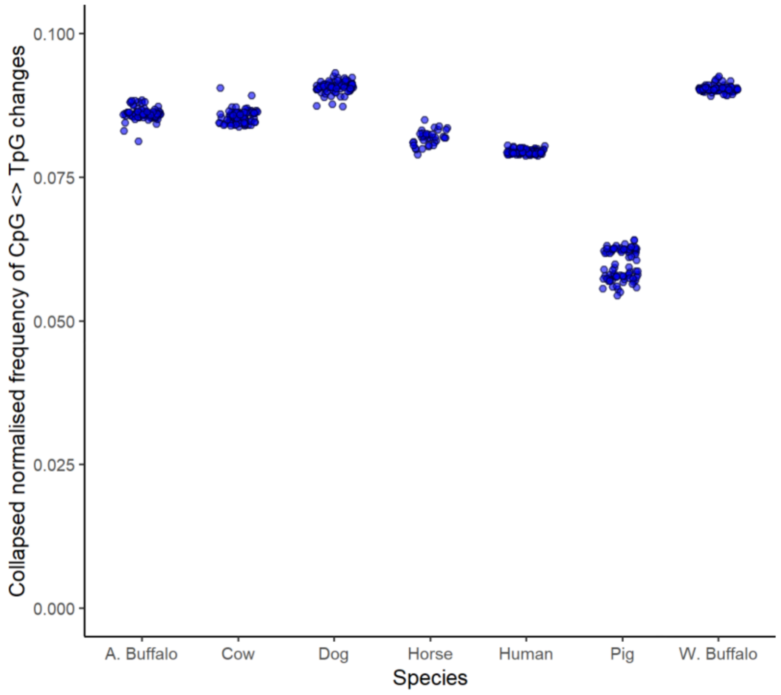


Supplementary Figure 9. The relative frequency of CpG<>TpG changes is lower in pigs, suggesting that the observed depletion of CpG>TpG changes in this species cannot simply be attributed to incorrectly inferring the direction of change at these sites. The split between the Asian and European pigs is also evident.

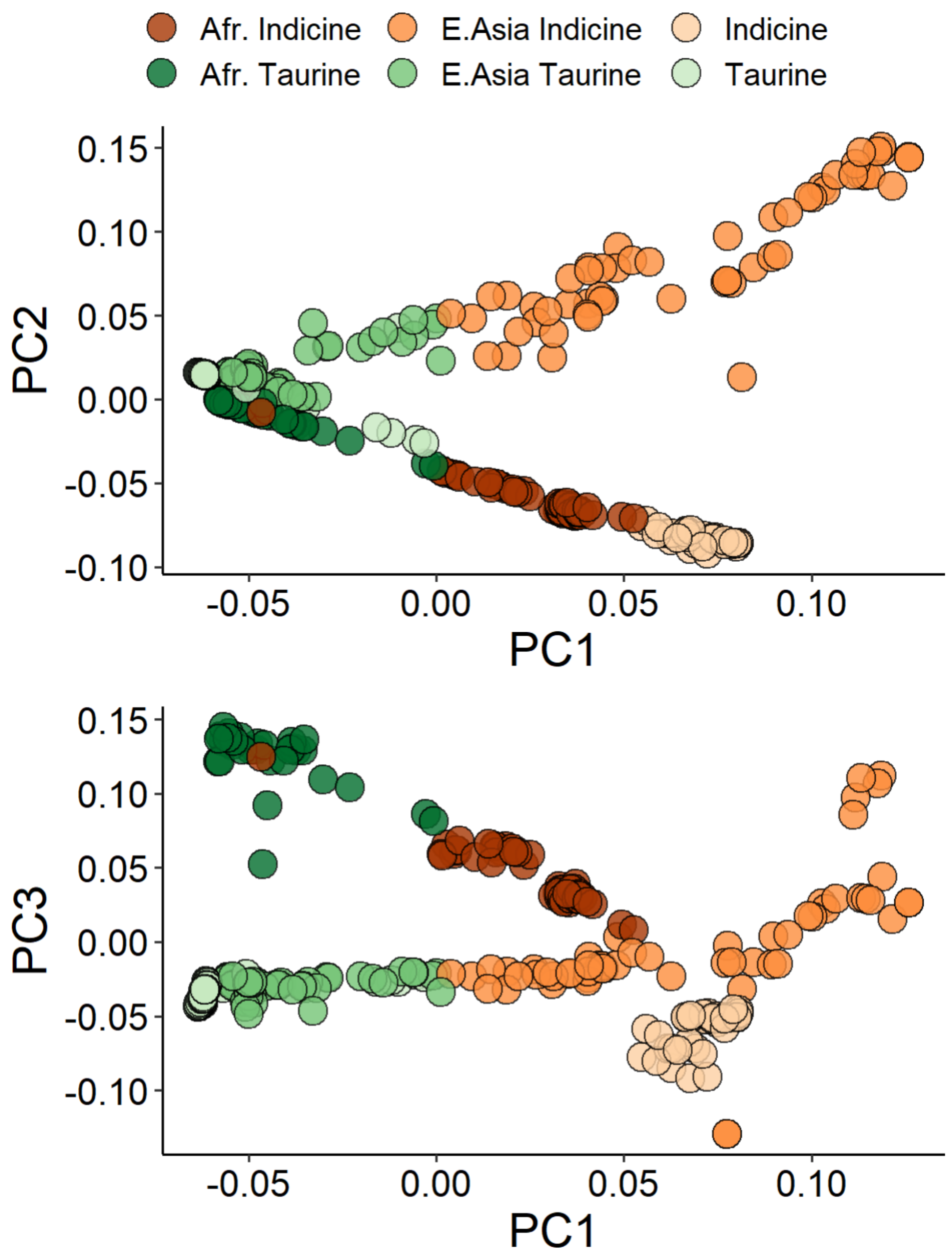


Supplementary Figure 10. Cattle PCA plots based on SNP genotypes (not mutation spectra).

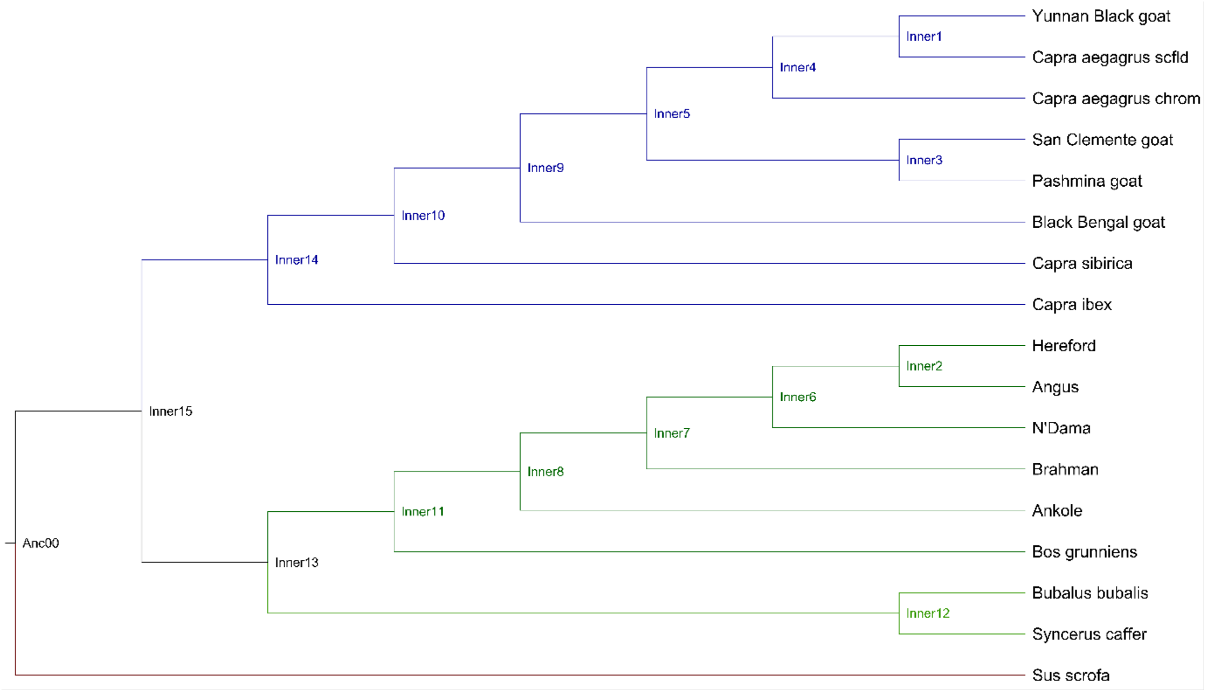


Supplementary Figure 11. The genomes included in the novel progressive cactus alignment in order to derive the bovid ancestral genomes.
